# Mesenchymal stromal cells release CXCL1/2/8 and induce chemoresistance and macrophage polarization

**DOI:** 10.1101/482513

**Authors:** Augustin Le Naour, Mélissa Prat, Benoît Thibault, Renaud Mével, Léa Lemaitre, Hélène Leray, Muriel Golzio, Lise Lefevre, Eliane Mery, Alejandra Martinez, Gwénaël Ferron, Jean-Pierre Delord, Agnès Coste, Bettina Couderc

## Abstract

Factors released by surrounding cells such as cancer-associated mesenchymal stromal cells (CA-MSCs) are involved in tumor progression and chemoresistance. We determine the mechanisms by which a naïve MSC could become a CA-MSC and characterize CA-MSCs. Ovarian tumor cells (OTC) trigger the transformation of MSCs to CA-MSCs expressing different pro-tumoral, genes and secreting high amounts of CXCR1/2 ligands (CXCL1, CXCL2 and IL-8) implicated in the chemoresistance of cancer cells. CXCR1/2 ligands can also inhibit the immune response against OTC. Indeed, through their released factors, CA-MSCs can trigger the differentiation of monocytes to pro-tumoral M2 phenotype macrophages known to promote the tumor progression. When CXCR1/2 receptors are inhibited, these CA-MSC-activated macrophages lose their M2 functions and acquire an anti-tumoral phenotype. Both *ex vivo* and *in vivo* a CXCR1/2 inhibitor can sensitize OTC to carboplatin even in the presence of a pro-tumoral microenvironment. This inhibitor can circumvent the pro-tumoral effects of CA-MSCs. As high concentrations of CXCR1/2 ligands in blood from patients can be associated with chemoresistance, CXCR1/2 inhibition could be a potential therapeutic strategy to revert chemoresistance.

## INTRODUCTION

Chemoresistance is a major problem concerning the treatment of cancers, whether they are solid or hematologic. Moreover, concerning ovarian cancers, resistance has been found to develop against many chemotherapeutic drugs. It can occur during treatment or several months post-treatment and is associated with an impaired prognosis. The chemoresistance of tumor cells could be caused by molecular alterations affecting metabolism, growth control and apoptosis pathways, uptake or efflux of the drug, targeted receptors, *etc*[1]. Chemotherapeutic agents may induce a rapid host response involving a “storm” of cells, cytokines and growth factors that promote angiogenesis, tumor regrowth, metastasis and chemoresistance[2–4]. Thus, chemoresistance could also be due to the microenvironment surrounding tumor cells.

The tumor microenvironment can include various different types of cells (endothelial cells, fibroblasts, adipocytes, immune cells, mesenchymal stromal cells (MSCs), *etc*.) as well as the extracellular matrix. In this study, we paid particular attention to MSCs.

MSCs are multipotent stromal cells that can differentiate into adipocytes, chondrocytes, osteoblasts, fibroblasts and vascular structures[5]. They can be isolated from different tissues including, among others, bone marrow, skeletal muscle, adipose tissue, the circulatory system[6] and tumors. In this last case, they are known as cancer-associated mesenchymal stromal cells (CA-MSCs)[7]. These cells have pro-tumoral properties and can stimulate growth and angiogenesis as well as the chemoresistance of tumor cells. This phenomenon occurs through the direct interactions of CA-MSCs with tumor cells[8] and/or the release of various factors including cytokines[9,10], growth factors[11], exosomes[12] and fatty acids[13] (for a review see Le Naour and Couderc[14]). Using a model of human ovarian adenocarcinoma, we previously described how factors released by CA-MSCs could be associated with chemoresistance. These factors activate the PI3K/Akt/mTOR pathway in ovarian cancer cells (OCT) to stabilize X-linked inhibitor of apoptosis protein (XIAP) and induce chemoresistance to carboplatin [15].

Concerning factors released by MSCs that could be involved in the acquisition of chemoresistance by tumor
cells, several authors have identified CCL5, IL-6 and the C-X-C motif chemokine ligand 8 (CXCL8 or IL-8)[16–18]. This last cytokine has been implicated in a number of inflammatory diseases involving the chemo-attraction and activation of neutrophils. Wang *et al.* have shown that the autocrine secretion of IL-8 by tumor cells induced their chemoresistance via a downregulation of the expression of effective caspases and the upregulation of several genes belonging to the multidrug resistance gene family such as MDR-1 or the anti-apoptotic gene family such as BCL-2 or XIAP[18]. The inhibition of IL-8 expression induces the chemosensitization of tumor cells. IL-8 and more extensively chemokines that bind to the same receptors, such as CXCR1 and CXCR2 ligands, could be involved in the acquisition of chemoresistance via the recruitment of MSCs around the tumor[18].

The signaling pathways activated by IL-8 (PI3K and phospholipase C) are stimulated though the interaction of the cytokine with CXCR1 and CXCR2 (CXCR1/2) (IL-8 receptor A and IL-8 receptor B) which are expressed by neutrophils, monocytes, endothelial cells, astrocytes, microglia and different types of tumor cells[19–21]. Browne *et al.* have described a strong correlation between the expression of CXCR1/2 and the grade of ovarian tumors [22].

Beside its role in chemoresistance, IL-8 can also stimulate the immune system[19]. It is a chemo-attractant for neutrophils, and could also interact with monocytes, as they express CXCR1/2. Monocytes differentiate into macrophages when they infiltrate tissues and represent then an important component of the ovarian tumor microenvironment (Tumor Associated Macrophages or TAM). They have great plasticity and can differentiate into several functional states in response to signals from the microenvironment. M1-macrophages have tumoricidal activity through the secretion of cytotoxic factors, while M2-macrophages generally only produce low levels of reactive nitrogen/oxygen species (ROS), exhibit low amounts of antigen-presentation and suppress anti-tumor immunity. Several studies have reported the recruitment of M2-macrophages to solid tumors in response to chemotherapy[23]. These macrophages provide an immunosuppressive microenvironment[24,25], participate in angiogenesis through the release of vascular endothelial growth factors (VEGF) and protect tumor cells against paclitaxel chemotherapy for solid tumors[26] or melphalan-induced apoptosis in the case of multiple myeloma[2].

The complex interactions between MSCs, macrophages and tumor cells involved in tumor growth, neo-angiogenesis and the acquisition of chemoresistance need to be elucidated, as the tumor environment ultimately determines the clinical behavior of the disease (progression and chemoresistance) and has a direct impact on the overall survival[27]. Ovarian cancer patients frequently form ascites, which refers to abnormal accumulation fluid in the peritoneal cavity. The fluid from which contains tumor cells, stromal cells and the factors secreted by these different cells.

In the present work, we isolated and characterized MSCs directly from biopsies of ovarian cancers to compare them to bone marrow – MSC (BM-MSCs). Then, we designed a model mimicking what could happen in the case of cancer and observed that malignant cells modify the phenotype of MSCs and induce their differentiation into CA-MSCs. We compared these “induced” CA-MSCs (iCA-MSC) to native BM-MSCs in terms of their potential to induce chemoresistance in ovarian tumor cells (OTCs) via the release of secreted factors, and moreover we identified CXCR1/2 ligands that could mediate the actions of CA-MSCs. As these receptors are also expressed on macrophages, we evaluated the effects of IL-8, CXCL1 and CXCL2, secreted by CA-MSCs, on the polarization of macrophages, in addition to their effects on the acquisition of chemoresistance by OTCs. Finally, we observed that CXCR1/2 inhibition could sensitize OTCs to chemotherapy and repolarize macrophages to an anti-tumoral phenotype.

We show that, in part, OTC chemoresistance appears to be due to CA-MSCs, which are found in ovarian tumors and are capable of secreting chemo-protective factors, including CXCL1, CXCL2 and IL-8 chemokines. High concentrations of chemokines in the serum of patients at the time of diagnosis is associated with carboplatin resistance. The combination of a CXCR1/2 inhibitor with conventional chemotherapy may be a promising therapeutic strategy to avoid resistance. In addition, this treatment could also be considered as an immunotherapy able to repolarize macrophages to an anti-tumoral phenotype.

## RESULTS

### 1. CA-MSCs isolated from tumor biopsies confer chemoresistance to OTCs

We isolated cells from freshly extracted human ovarian adenocarcinoma biopsies (n=12). These cells were selected based on their adherence to plastic dishes. They presented a fibroblast-like morphology as BM-MSC (Figure 1A and 1B). These cells were then sorted according to the expression of CD73, CD90 and CD105 (Figure 1C), which are classic markers of MSC. They present the same phenotype as BM-MSC as they express CD73, CD90 and CD105 and do not express CD14, CD20, CD34 and CD45. These cells are “cancer-associated mesenchymal stromal cells” (CA-MSCs).

**Figure 1:**
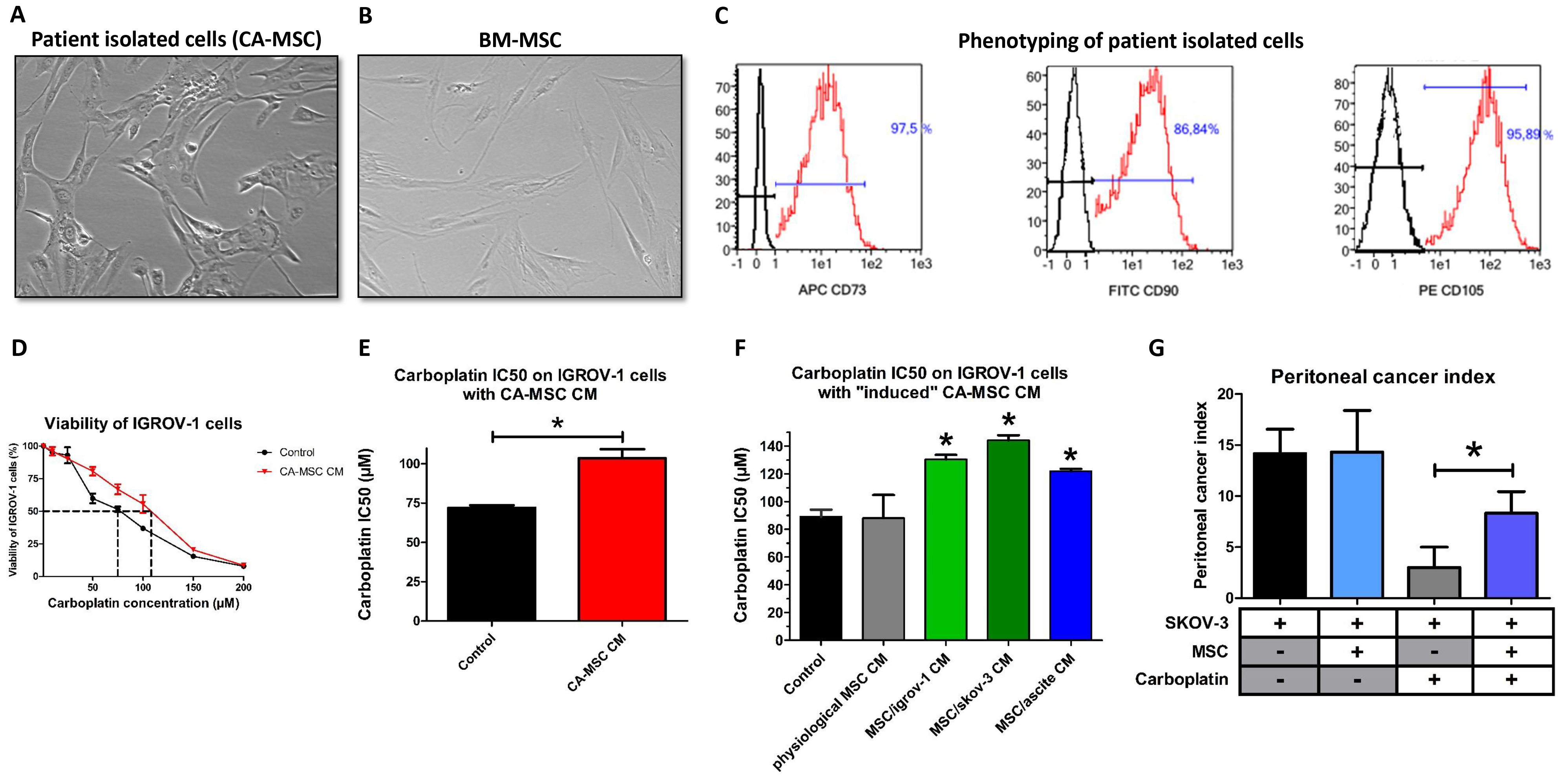
The acquisition of chemoresistance by ovarian tumor cells (OTCs) through factors secreted by Carcinoma associated-mesenchymal stromal cells (CA-MSCs) A, B: Phenotype of stromal cells from patient biopsies and bone marrow-mesenchymal stromal cells (BM-MSCs) (X40). C: Flow cytometry analysis of stromal cells from patient biopsies. The expression of CD73, CD90 and CD105 was evaluated. D: CA-MSCs protect OTCs from carboplatin-induced growth inhibition. OTCs cultured alone or in the presence of CA-MSC CM were treated with increasing concentrations of carboplatin for 48 hours. Cell viability was measured for IGROV-1 cells cultured in control medium or in CA-MSC conditioned medium (CM). The dotted line corresponds to 50% cell viability. E: Histogram representing the mean IC50 of carboplatin on IGROV-1 cells cultured with CA-MSC CM (n=12). F: Histogram representing the IC50 for carboplatin on IGROV-1 cells cultured with BM-MSC CM (physiological MSCs) or “induced” CA-MSC (iCA-MSC) CM from different origins (BM-MSCs cultivated with IGROV-1 CM, SKOV-3 CM or ascites) (n=4 for each type of MSC). G: The mean value of the peritoneal cancer index +/− SEM is presented for the mice injected with SKOV-3 cells with and without MSCs and treated (+) or not (−) with carboplatin (n=6 mice/group of treatments). The comparison between groups was performed using a Wilcoxon-Mann Whitney test (independent non-parametric data). P values of <0.05 (*) indicate a significant difference.

Next, we evaluated the ability of CA-MSCs to induce chemoresistance in OTCs. For this purpose, we cultured IGROV-1 cells, a human OTC line, in the presence of conditioned media (CM) from CA-MSCs and treated the cells with carboplatin, which is the gold standard in ovarian cancer. Cell viability was evaluated after 48 hours and the concentration of carboplatin that inhibited 50% of cell viability (IC50) was calculated (Figure 1D). As shown Figure 1E, CA-MSC CM induced an increase of 44% in the IC50 of carboplatin on IGROV-1 cells. All the CA-MSCs that we cultured (n=12) could induce chemoresistance through released factors (Figure 1E and S1A). CA-MSC CM also induced an increase of 33% in the IC50 of carboplatin on another OTC line (SKOV-3 cells) (Figure S1B).

### 2. BM-MSCs could differentiate into CA-MSCs in a tumoral microenvironment

MSCs can display different phenotypes and functions, depending on the type of tissue from where they are isolated, including, among others, ovary, bone marrow, adipose tissue, heart and bladder [28,29]. Thus our aim was to analyze if CA-MSCs acquired specific functions because of their surrounding tumor cells. We hypothesized that CA-MSCs isolated from ovarian nodules could be resident MSCs related to BM-MSCs, which may have been recruited to the tumor because of the inflammatory context and the chemokines released by tumor cells. Those recruited MSCs could have been educated by tumor cells to adopt new functions such as the ability to induce chemoresistance. To find out if CA-MSCs could be differentiated cells derived from progenitor MSCs, we cultured multipotent BM-MSCs from healthy female donors for 21 days either in control medium (complete DMEM) (physiological BM-MSCs) or in CM obtained from two different human OTC lines (IGROV-1 or SKOV-3) or in ascitic fluid (ascites) from patient (here we use ascites in which cells have been removed by centrifugation and filtration).

After 21 days of culture in the different conditions, the BM-MSC media was changed for complete DMEM. After three days, these media were isolated and called “physiological MSC CM” (when the BM-MSCs had been cultured in control medium) or “induced CA-MSC (iCA-MSC) CM” when the BM-MSCs had been cultured in OTC CM and ascites. Indeed iCA-MSCs were differentiated from BM-MSCs which have been cultured in IGROV-1 CM (MSC/igrov-1), SKOV-3 CM (MSC/skov-3) or ascitic fluid (MSC/ascite CM). Their media were tested for their ability to induce chemoresistance in IGROV-1 cells. As shown Figure 1F, while the physiological MSC CM did not confer chemoresistance to OTCs, iCA-MSC CM induced an increase in the chemoresistance of IGROV-1 cells to carboplatin, similar to that observed with CA-MSC CM (Figure 1E and S1A). By culturing BM-MSCs in a tumoral context (secreted factors from OTCs or ascites) we modified their capabilities, they gained the ability to secrete factors that confer chemoresistance to OTCs, as was observed with tumor extracted CA-MSCs.

In order to verify our hypothesis *in vivo*, we injected ovarian tumor bearing nude mice (i.p. injection of 10 × 10^6^ human SKOV-3 cells) with 1 × 10^6^ BM-MSCs from healthy donors and analyzed their ability to confer chemoresistance to OTCs. SKOV-3 cells have been chosen because of the reproducible implantation of a tumor into animals using these cells compared to the IGROV-1 cells. Mice were injected with carboplatin once a week for three weeks beginning 10 days post BM-MSC injection. We waited 10 days before beginning the treatment, in order to allow the OTCs to interact with and potentially activate the BM-MSCs so to be as close as possible to the *ex vivo* experiment previously shown. We evaluated tumor progression seven days after the last injection of carboplatin by measuring the peritoneal cancer index (Table 1) as previously described [30]. Briefly, we analyzed the number of nodules and the size of the biggest nodule (as depicted in Table 1) and calculated the peritoneal cancer index. The injection of BM-MSCs into tumor bearing mice (SKOV-3) did not modify tumor progression (Figure 1G) but did modify the efficiency of the carboplatin treatment (chemoresistance). BM-MSCs injected into mice at the same time as OTCs conferred chemoresistance to OTCs *in vivo* (Figure 1G). BM-MSCs in the vicinity of OTCs could acquire a CA-MSC phenotype which in turn induced the ovarian cancer cell chemoresistance that we noticed *in vivo*.

**Table 1:**
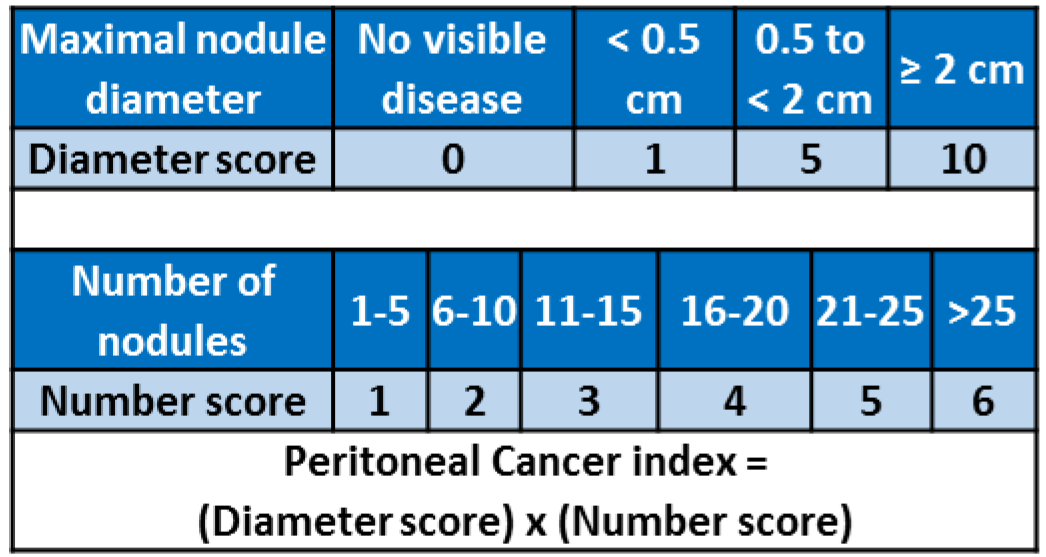
Peritoneal cancer index. Table outlining the parameters used to evaluate the peritoneal cancer index.

### 3. CA-MSCs and iCA-MSCs lose their multipotency

We evaluated whether any of the three major signaling pathways involved in chemoresistance were activated in MSCs cultured in OTC CM. As shown Figure 2A and 2B, the PI3K/Akt, MAPK and NF-κB signaling pathways were activated in iCA-MSCs and this was observed either in iCA-MSCs derived from MSCs cultured in CM from OTC lines (IGROV-1).

**Figure 2:**
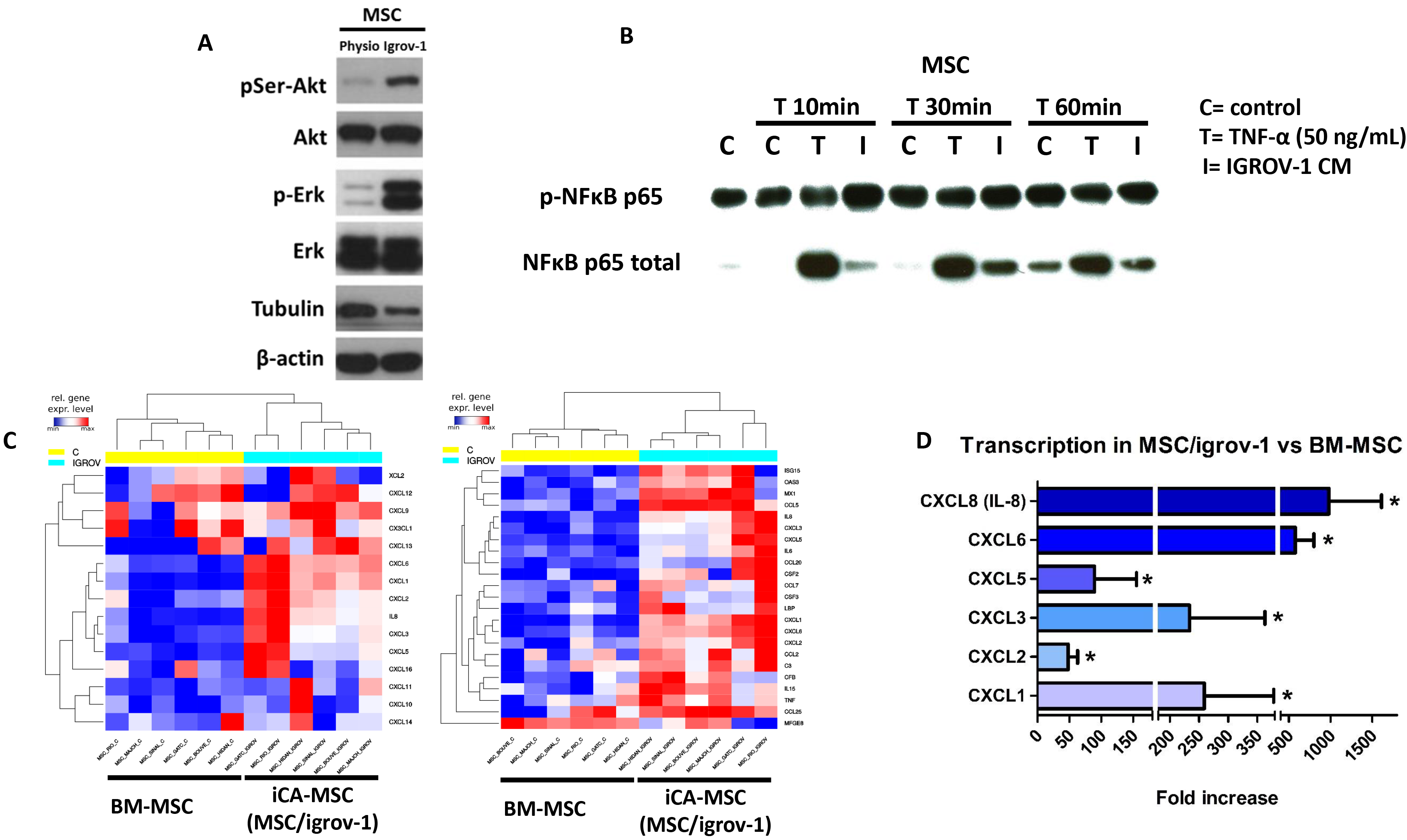
Factors secreted by OTCs activate PI3K/Akt, MAPK and NF-κB signaling pathways and modify gene expression in CA-MSCs. A: BM-MSCs were cultured in IGROV-1 CM or in control medium (physiological BM-MSCs) for 21 days. Cells were collected and proteins were extracted. Akt, phospho-Akt (Ser 473), Erk and Phospho-Erk expression was assayed by western blot. Phospho-Akt expression was normalized to total Akt expression. Phospho-Erk expression was normalized to total Erk expression (mean+/−SEM, n=3). B: BM-MSCs were cultured with IGROV-1 CM or with control medium (control) for the indicated amount of time. T represents treatment of the BM-MSCs with 50ng/mL of TNFα. Cells were collected and proteins were extracted. NF-κB and phospho-NF-κB p65 expression was assayed by western blot. C: Representative gene expression of the two type of cells (physiological BM-MSCs and iCA-MSCs (MSC/igrov-1). BM-MSCs were cultured for 21 days either in control media (MSC/physio) (n=6) or in the presence of IGROV-1 CM (MSC/igrov-1 = iCA-MSC) (n=6), RNA was extracted and analyzed by Nanostring™ technology D: quantification of the transcriptomic analysis showing the fold induction of the selected genes. Transcriptomic analysis performed using Nanostring technology: analysis of all the genes (n = 770) found on the chip[31], required an initial filter (sd>0.25) to eliminate genes with little variation. An ANOVA test was applied to compare the six BM-MSC control samples versus the six iCA-MSC (MSC/igrov-1) samples. The p values were corrected for multiple tests using the Benjamini & Hochberg (BH) method.

As BM-MSCs are multipotent cells, we evaluated whether CA-MSCs (isolated from patient tumors) were able to differentiate into osteoblasts or adipocytes. We used BM-MSC as control of differentiation and observed that BM-MSCs were able to differentiate into either osteoblasts or adipocytes. CA-MSCs were found to not be multipotent, as they had lost their ability to differentiate into osteoblasts or adipocytes (Figure S2A and S3A). BM-MSCs that were cultured for 21 days in control medium were still multipotent. On the contrary, BM-MSCs cultured in a tumoral environment lost their multipotency as when we tested the multipotency of the iCA-MSCs (Figure S2B and S3B), these cells were found to not be multipotent. Tumoral secreted factors induced the differentiation of multipotent BM-MSCs into iCA-MSCs, which lost their multipotency. CA-MSC seemed not to be cancer associated fibroblasts are derived from MSCs as they do not express the αSMA and FAP expression

In these experiments, we found that when we culture BM-MSCs in OTC CM they acquire additional functions such as the ability to secrete factors that induce chemoresistance in OTCs, in addition they lose their multipotency. We observed a crosstalk between MSCs and OTCs. OTCs can activate MSCs, which are able in turn to help OTCs become resistant to chemotherapy.

### 4. CA-MSCs and iCA-MSCs overexpress the chemokines CXCL1, CXCL2 and IL-8

We observed that iCA-MSCs acquired functions such as the ability to secrete factors able to induce chemoresistance in OTCs, this was due to their tumoral environment and perhaps due to the activation of the PI3K/Akt, MAPK and NF-κB signaling pathways (Figure 2A and 2B). In order to compare gene expression in BM-MSCs and their derived iCA-MSCs, we used the nCounter^®^ Analysis System[31] to investigate the genes differentially expressed between iCA-MSCs (MSC/igrov-1) and physiological BM-MSCs in six pairs of samples. With this method, it is possible to compare BM-MSCs and iCA-MSCs from the same donor and overcome interindividual variability. Each pair corresponds to BM-MSCs cultured for 21 days in control medium or in IGROV-1 CM. As shown Figure 2C and D, iCA-MSCs overexpressed several pro-tumoral genes. They also overexpressed pro-metastatic (such as CCL5), and pro-angiogenic genes (e.g. CXCL8 and CCL5). The modification in MSC gene expression that we observed confirms that culturing MSCs in a tumoral context can modify their phenotype to a pro-tumoral phenotype. Clearly MSCs are influenced by their environment.

To identify the secreted factor over produced by CA-MSCs that mediates the acquisition of chemoresistance by tumor cells, we analyzed *in silico* data obtained from the gene expression analysis. We focused on secreted factors shown in the literature to be involved in the acquisition of chemoresistance and the activation of either the PI3K/Akt or the NF-κB signaling pathways, as previously described [10]. Between the candidates, we retained chemokines and more specifically the CXCR1/2 ligands as CXCL1 (fold increase = 259), CXCL2 (fold increase = 48) and IL-8 (fold increase = 979), that were overexpressed in iCA-MSCs compared to BM-MSCs (Figure 2D) and these chemokines are known to be involved in the chemoresistance of OTCs to cisplatin and taxan[18] or more generally in solid tumor progression [32].

In order to confirm the results obtained by the nCounter^®^ analysis, we performed RT-qPCR analysis on BM-MSCs (cultured for 21 days in control medium (physiological)) or their counterpart iCA-MSCs (obtained after BM-MSCs were cultured for 21 days in IGROV-1 CM, SKOV-3 CM or ascites) and compared the expression of CXCL1, CXCL2 and IL-8. As shown Figure 3A, 3B and 3C, there was a strong overexpression of CXCL1, CXCL2 and IL-8 mRNA in all three types of iCA-MSCs compared to the control.

**Figure 3:**
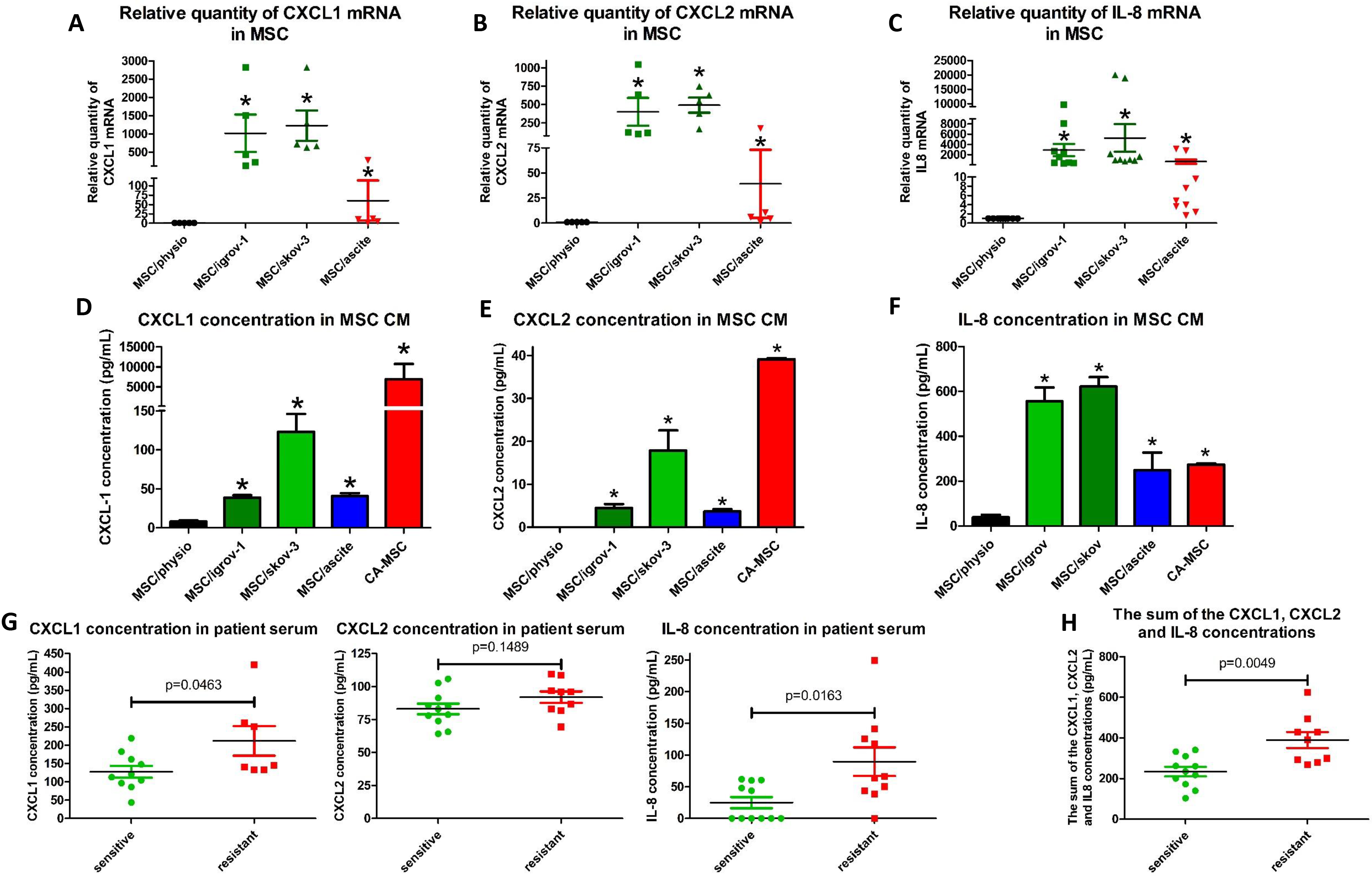
Determination of the CXCR1/2 ligands secreted by CA-MSCs and iCA-MSCs. BM-MSCs were cultured alone or in CM from IGROV-1 (MSC/igrov-1) or SKOV-3 (MSC/skov-3) cells or ascites (MSC/ascite) for 21 days. Then media were changed for complete DMEM. After three days cells or CM were collected. A-C: The upregulation of genes coding for CXCL1, CXCL2 and IL-8 was validated by RT-qPCR performed on RNA extracted from BM-MSCs and different types of iCA-MSCs (induced by IGROV-1 CM, SKOV-3 CM or ascites).The data from BM-MSCs were set to 1 and the relative quantity of mRNA is shown. CXCL1 (A), CXCL2 (B), IL-8 (C). D - F: The concentrations of CXCL1, CXCL2 and IL-8 in the CM were quantified using an ELISA kit. The CM from CA-MSCs was also tested. Histograms show the mean concentrations of three independent experiments performed in triplicate. (Mean +/− SEM, * p < 0.05). CXCL1 (D), CXCL2 (E), IL-8 (F). G Determination of the concentration of CXCR1/2 ligands in patient samples (n=21): The concentrations of CXCL1, CXCL2 and IL-8 were determined using ELISA kits on samples of serum from patients with ovarian adenocarcinoma collected at diagnosis. Tumors were given the status of “sensitive” (n=11 patients) or “resistant” (n = 10 patients), determined *a posteriori* according to the duration of the platinum-free interval before relapse. H: The sum of the concentrations of CXCL1, CXCL2 and IL-8 was obtained by adding together the serum concentration of these three chemokines. The comparison between groups was performed using a Wilcoxon-Mann Whitney test (independent non-parametric data). P values of <0.05 (*) indicate a significant difference.

To confirm the RT-qPCR analysis, we evaluated the concentration of the cytokines CXCL1 (Figure 3D), CXCL2 (Figure 3E) and IL-8 (Figure 3F) in the CMs from CA-MSCs isolated from patients or from the different iCA-MSCs compared to CM from BM-MSCs. We observed a much higher concentration of the three chemokines in the CM from CA-MSCs or iCA-MSCs compared to that observed in the CM from BM-MSCs.

Finally, we evaluated the amount of the three CXCR1/2 ligands in serum from patients with ovarian adenocarcinoma collected at diagnosis. Tumors were given the status of “sensitive” or “resistant” as determined *a posteriori*. The classification of the relapse is obtained according to the duration of the platinum-free interval, corresponding to the time between the date of the last dose of platinum and the date of the relapse. Cancers whose recurrence is longer than 6 months are considered sensitive to platinum and if it is less than 6 months, it is considered resistant [33]. Table 2 presents the characteristics of the 21 patients. As shown Figure 3G, there was an increased concentration of the three chemokines in the serum from patients with resistant tumors compared to those with sensitive tumors with p values of 0.046, 0.149 and 0.016 for CXCL1, CXCL2 and IL-8, respectively. The sum of the concentrations of CXCL1, CXCL2 and IL-8 was obtained by adding together the serum concentrations of these three chemokines, and it showed that patients with resistant tumors had a higher serum concentration of these three chemokines (p = 0.0049). Our results showed that increased levels of CXCR1/2 ligands (CXCL1, CXCL2 and IL-8) could be associated with future tumor resistance to chemotherapy and may be predictive markers of a tumor’s response to chemotherapy. Thus, they are a good target to explain the mechanism by which MSCs induce the acquisition of chemoresistance in OTCs.

**Table 2:** Baseline characteristics of patient samples.

| Number | Type | Stage | Chemotherapy | Free interval | Sensitivity |
| --- | --- | --- | --- | --- | --- |
| 1 | High-grade serous ovarian carcinoma | IIIc | Carboplatin/paclitaxel | >36 months | Sensitive |
| 2 | Ovarian carcinoma with low differentiation | IV | Carboplatin/paclitaxel | Progressive disease | Refractory |
| 3 | High-grade serous ovarian carcinoma | III | Carboplatin/paclitaxel | 30 months | Sensitive |
| 4 | Serous papillary ovarian carcinoma | IIIc | Carboplatin/paclitaxel | 5 months | Resistant |
| 5 | Serous papillary ovarian carcinoma | IV | Carboplatin/paclitaxel | >48 months | Sensitive |
| 6 | Clear cell carcinoma | IIIc | Carboplatin/paclitaxel | 7 months | Sensitive |
| 7 | High-grade serous ovarian carcinoma | II | Carboplatin/paclitaxel | 14 months | Sensitive |
| 8 | Serous papillary ovarian carcinoma | IIIb | Carboplatin/paclitaxel | 30 months | Sensitive |
| 9 | High-grade serous ovarian carcinoma | IIIc | Carboplatin/paclitaxel | 22 months | Sensitive |
| 10 | Serous papillary ovarian carcinoma low grade | III | Carboplatin/paclitaxel | >60 months | Sensitive |
| 11 | Atypical mucinous carcinoma | III | Carboplatin/paclitaxel | 3 months | Resistant |
| 12 | High-grade serous ovarian carcinoma | IV | Carboplatin/paclitaxel | 8 months | Sensitive |
| 13 | High-grade serous ovarian carcinoma | III | Carboplatin/paclitaxel | Progressive disease | Refractory |
| 14 | Serous ovarian carcinoma | IIIc | Carboplatin/paclitaxel | >42 months | Sensitive |
| 15 | Low-grade serous ovarian carcinoma | III | Carboplatin/paclitaxel | 18 months | Sensitive |
| 16 | Serous ovarian carcinoma | III | Carboplatin/paclitaxel | 4 months | Resistant |
| 17 | High-grade serous ovarian carcinoma | IV | Carboplatin/paclitaxel | 3 months | Resistant |
| 18 | Serous ovarian carcinoma with moderate differentiation | ND | Carboplatin/paclitaxel | Progressive disease | Refractory |
| 19 | Serous ovarian carcinoma | IV | Carboplatin/paclitaxel | 3 months | Resistant |
| 20 | High-grade serous ovarian carcinoma | III | Carboplatin/paclitaxel | 5 months | Resistant |
| 21 | Ovarian adenocarcinoma with low differentiation | IV | Carboplatin/paclitaxel | Progressive disease | Refractory |

**Table 3:**
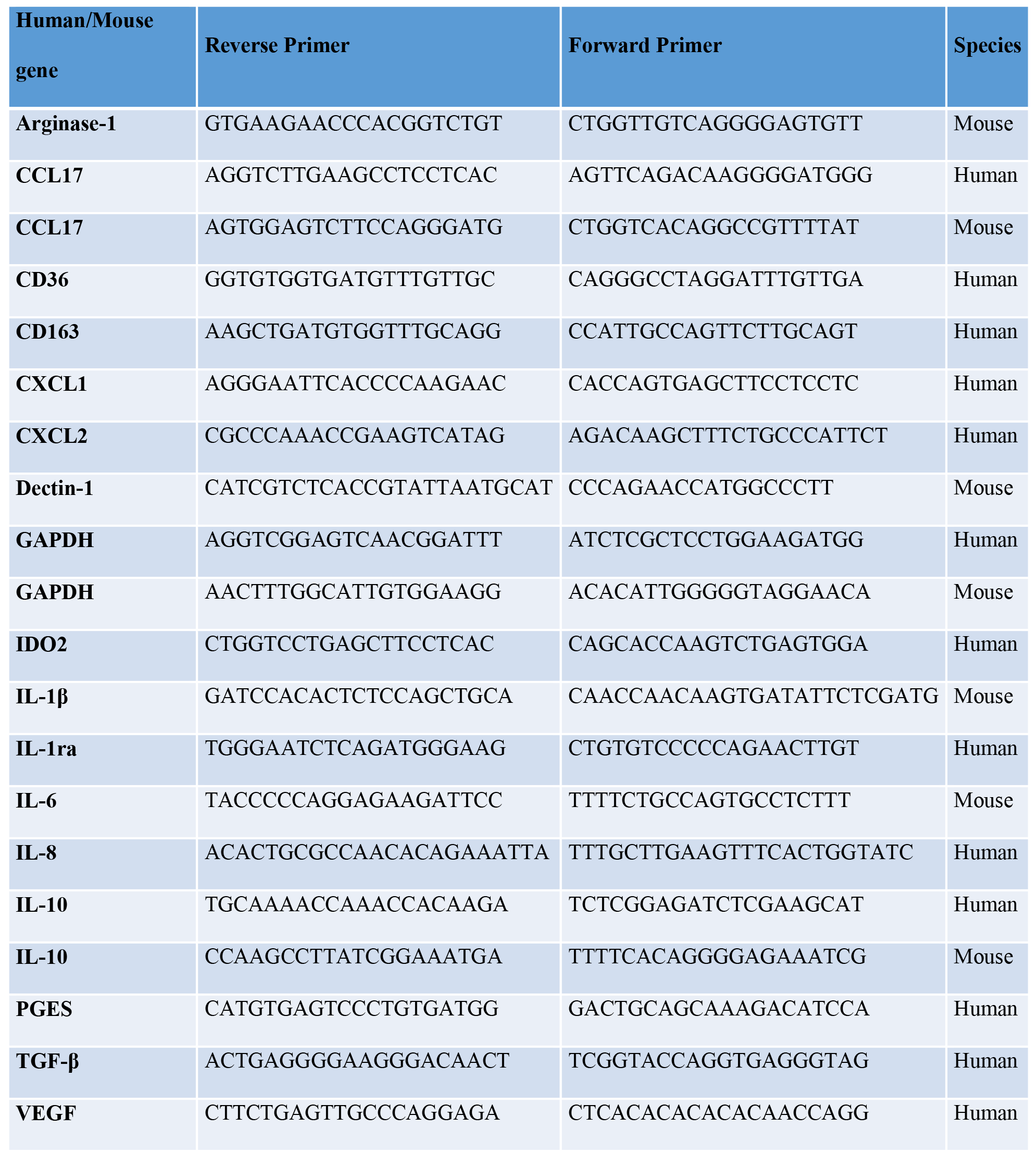
Primers used for qRT-PCR analysis.

### 5. CXCR1/2 inhibition and chemoresistance

In order to determine whether CXCR1/2 ligands could play a role in the chemoresistance mediated by MSCs, we determined the IC50 of carboplatin on IGROV-1 cells in the presence of an inhibitor directed against CXCR1/2 receptors[34]. We previously verified that CXCR1/2 were expressed in different human OTC lines and showed that epithelial adenocarcinoma cells lines (OVCAR-3, IGROV-1 and SKOV-3 cells) as well as a clear cell carcinoma cell line (JHOC-5) expressed CXCR1 as well as CXCR2 (Figure S4A and S4B).

While the CXCR1/2 inhibitor did not alter the viability of tumor cells alone (Figure S4C), it induced an increase in their sensitivity to carboplatin when they were cultured in complete medium alone. This is due to the inhibition of the autocrine production of IL-8 by the tumor cells [9]. The acquisition of chemoresistance by IGROV-1 cells mediated through factors secreted by CA-MSCs or iCA-MSC could be reversed by the presence of the CXCR1/2 inhibitor (Figure 4A). The acquisition of chemoresistance by IGROV-1 cells mediated through factors secreted by iCA-MSCs could also be reversed with the CXCR1/2 inhibitor (Figure 4B). The CXCR1/2 inhibitor was able to revert the chemoresistance acquisition mediated by CA-MSCs as well as sensitive OTCs to carboplatin.

**Figure 4:**
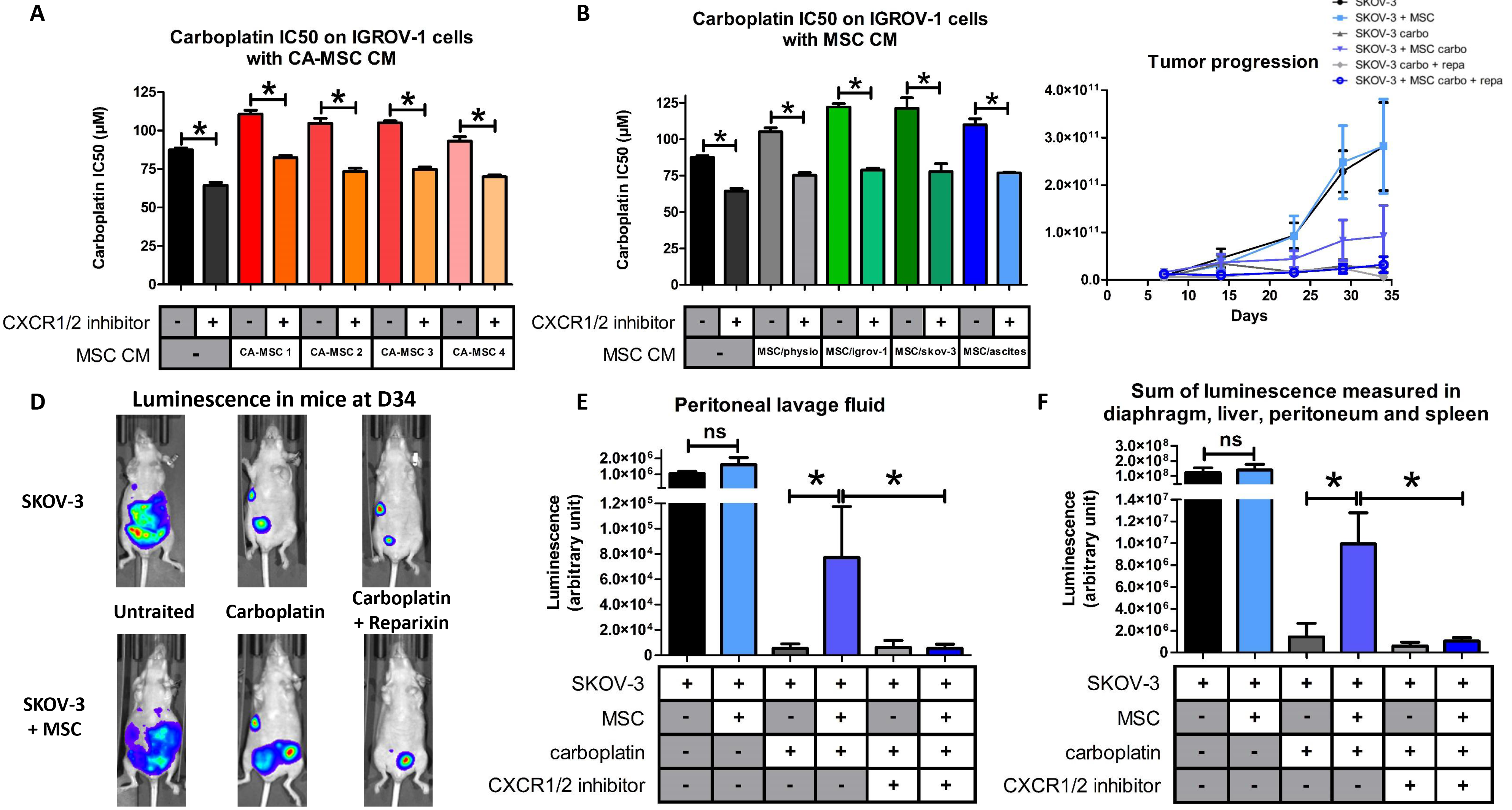
Implication of the CXCR1/2 axis in OTC resistance to carboplatin. A, B: The IC50 of carboplatin was monitored as described previously in Figure 1. IGROV-1 cells were cultured in the presence or not of CM from BM-MSCs (MSC/physio), CA-MSCs (n=4) or iCA-MSCs (MSC/igrov-1 and MSC/skov-MSC/ascites). At day 1, cells were treated with carboplatin admixed or not with a CXCR1/2 inhibitor (100 μM). Cell viability was evaluated at day 3. C: 10^7^ SKOV-3luc +/− 10^6^ MSC were injected intraperitoneally (i.p.) into nude mice (8 mice/group at the beginning of the experiments). After 7 days, treatments were started, including 1 injection of carboplatin every 7 days for 3 weeks and 1 injection of reparixin 3 times per week for 3 weeks. Analysis of bioluminescence through the whole body of the mice was performed once a week after an i.p. injection of luciferin. At day 34, bioluminescence of the whole body of the mice was analyzed making it possible to obtain photographs to vizualize and to quantify the luminescence illustrated here using one mouse per group. D, E: On day 36, the mice were euthanized and a peritoneal lavage was carried out with 5 mL of NaCl 0.9%. The peritoneum, spleen, liver and diaphragm were removed. After addition of luciferin, the sum of the luminescence of the peritoneum, spleen, diaphragm and liver (D) and the luminescence in the peritoneal lavage fluid (E) was measured. The comparison between groups was performed using a Wilcoxon-Mann Whitney test (independent non-parametric data). P values of <0.05 (*) indicate a significant difference.

We used an *in vivo* murine model to confirm that the CXCR1/2 inhibition could reverse the acquisition of chemoresistance by OTCs. We injected nude mice with ovarian adenocarcinoma cells (10.10^6^ human SKOV-3 cells) admixed or not with human MSCs (1.10^6^ cells). Mice were treated or not with carboplatin admixed or not with the CXCR1/2 inhibitor reparixin (usable *in vivo*) [35]. The progression of ovarian cancer was monitored by measuring bioluminescence both in real time at different time points post tumor injection, and in the different organs and peritoneal lavage fluid after the mice were sacrificed 36 days (D36) post tumor injection. As shown in Figure 4C, the co-administration of MSCs with OTCs did not modify tumor development. Treatment of mice with carboplatin reduced tumor development, unless the mice were co-administered with OTCs and MSCs. We confirmed here that MSCs induced chemoresistance in human OTCs. Reparixin treatment reversed the chemoresistance induced by MSCs, as tumor progression was abolished when mice were treated with a mix of carboplatin and reparixin (Figures 4D and 4E).

We confirmed that inhibition of CXCR1/2 can reverse the acquisition of chemoresistance mediated by factors released by MSCs (CXCR1/2 ligands).

### 6. MSCs could play a role in the anti-tumoral activity of immune cells

Several authors, including us, have shown that factors secreted by MSCs at the MSC-macrophage interface are involved in re-educating macrophages by manipulating metabolic programs in differentially polarized macrophages [36,37]. Some cytokines have been reported to play a role, including, in addition to IL-8 [38], IL-6 and LIF [37,39]. Transcriptomic analysis (Figure 2A) of iCA-MSCs (MSC/igrov-1) and normal BM-MSCs (six pairs of samples) revealed that iCA-MSCs overexpress factors involved in the activation of immune cells, in particular the CXCR1/2 ligands, IL-6 and LIF (whose upregulation was verified by RT-qPCR, see Figure S4D and S4E) and CCL5, CXCL3, CXCL5 and CXCL6. Macrophages are activated either classically (M1 phenotype) or alternatively (M2 phenotype) [40]. M1 macrophages are classically anti-tumoral, whereas M2 are pro-tumoral. M2-polarized macrophages express high levels of CD206, CD163 and TGFβR, whereas M1 macrophages express high levels of CD40, CD80 and CD86 on their cell surface. Ovarian cancer patients with a poor clinical outcome and chemoresistance have a high proportion of M2 polarized macrophages[41].

To clarify the role of factors secreted by MSCs in macrophage polarization, we cultured naive monocytes from healthy donors in the presence of CM from BM-MSCs (physiological), CA-MSCs or iCA-MSCs for two days and performed an RT-qPCR analysis. For each gene, as shown in Figure 5A, we compared the quantity of mRNA between monocytes/macrophages that were cultured in macrophage SFM (control medium) or in CM from BM-MSCs (MSC/physio) or iCA-MSCs (MSC/igrov-1 or MSC/skov-3). . The levels of expression of M2 specific genes (Figure 5A) were increased in the monocytes/macrophages cultured in the presence of CM from iCA-MSCs. There was no significant difference in the expression of any tested M1 specific genes (CD80, CD16, CD86 …) Therefore, iCA-MSC CM was able to induce the upregulation of M2 specific markers, suggesting a conversion of naive monocytes to M2 macrophages.

**Figure 5:**
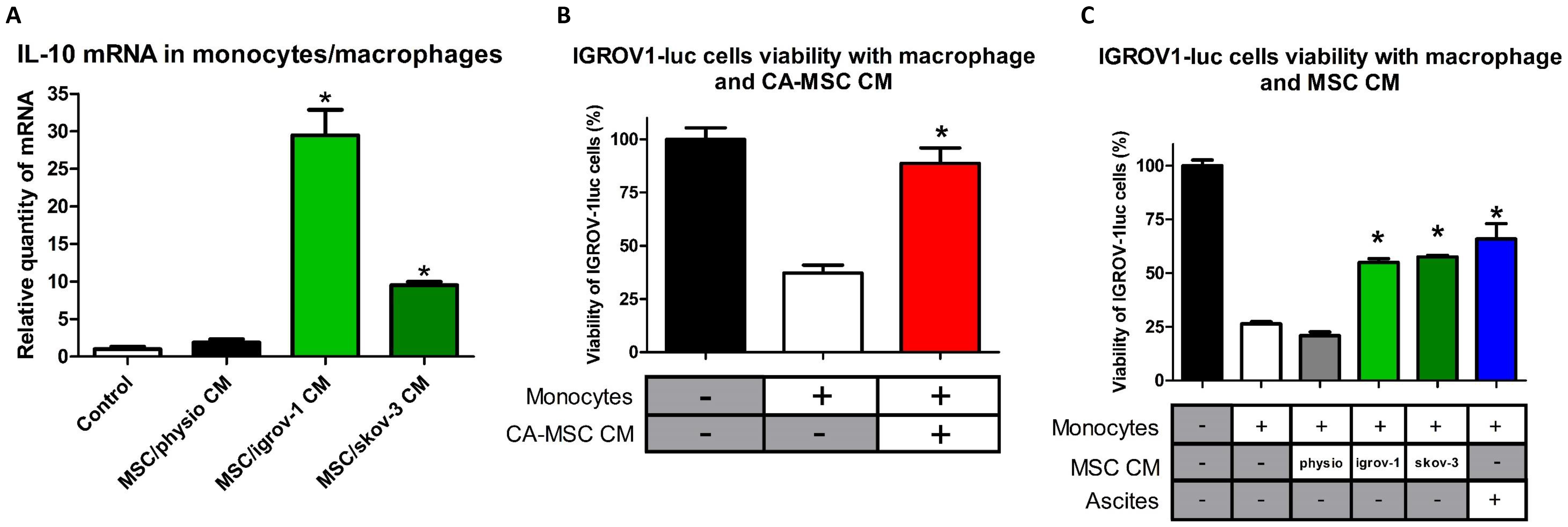
CA-MSCs interact with macrophages and facilitate monocyte to macrophage differentiation towards the TAM phenotype. A: Human monocytes were cultured for 2 days in control media, or in CM from BM-MSCs (MSC/physio CM) or in CM from different types of iCA-MSCs (induced by IGROV-1 CM (MSC/igrov-1 CM) or SKOV-3 CM (MSC/skov-3 CM). After culture, cells were washed and mRNA expression levels were analyzed by RT-qPCR (A). The data from control media were set at 1 and the relative quantity of mRNA is shown. B, C: Bar graphs representing the % of IGROV-1 cells viability monitored by measuring luciferase activity (n = 4). The cytotoxic activity of the macrophages that have been cultured in different media (CM of CA MSCs (left) or CM of BM-MSCs (physio) or of iCA-MSCs (MSC/igrov-1 CM or MSC/skov-3 CM) or ascites (right) on the IGROV-1luc cells was evaluated by measuring luciferase activity. The comparison between groups was performed using a Wilcoxon-Mann Whitney test (independent non-parametric data). P values of <0.05 (*) indicate a significant difference.

As M2 macrophages (M2b, M2c and M2d) are pro-tumoral and do not have tumoricidal functions, we analyzed the tumoricidal functions of these polarized macrophages. We cultured genetically modified IGROV-1 OTCs (IGROV1-luc) with monocytes that had been activated in control medium or in CM from CA-MSCs (Figure 5B) or from various iCA-MSCs (MSC/igrov-1 or MSC/skov-3) or in ascites (Figure 5C). Monocytes were cultured in ascites as a positive control for the M2 polarization of naïve monocytes, as has been previously described by Duluc *et al.* [39]. We evaluated the viability of the OTCs after three days by measuring luciferase activity. We observed that when the cells were co-cultured in control medium or in BM–MSC CM, the monocytes were able to kill between 70 to 80% of the OTCs. On the contrary, when the naïve monocytes were cultured in CM from CA-MSCs or iCA-MSCs they were not able to kill OTCs and allowed them to proliferate (Figures 5B and 5C).

In addition to their role in the acquisition of chemoresistance by OTCs, CA-MSCs could also be involved in the polarization of macrophages and the loss of their tumoricidal functions.

### 7. The tumoricidal function of macrophages could be restored by a CXCR1/2 inhibitor

As the CA-MSC and the iCA-MSC induced a modification of the tumoricidal activity of the macrophages unlike the BM-MSC and because this observation could be associated with a difference in the secretion of CXCL1 / 2 and IL-8 between these cells, we analyzed the impact of the CXCR1/2 inhibitor on the ability of macrophages cultured in CA-MSC CM to kill OTCs. We cultured genetically modified IGROV-1 OTCs (IGROV1-luc) with naïve monocytes in control medium or in CM from CA-MSCs or from various iCA-MSCs (MSC/igrov-1 and MSC/skov-3) or in ascites in the presence or absence of the CXCR1/2 inhibitor. We evaluated the viability of the OTCs after three days by measuring luciferase activity. As shown Figure 6A, we observed that while the naïve monocytes that had been cultured in CM from CA-MSCs or iCA-MSCs or ascites did not to kill effectively OTCs, the same monocytes cultured in the same conditions but in the presence of the CXCR1/2 inhibitor had tumoricidal properties. The effect of the CXCR1/2 inhibitor was more pronounced on the monocytes cultured in CM from CA-MSCs than on the monocytes cultured in ascites.

**Figure 6:**
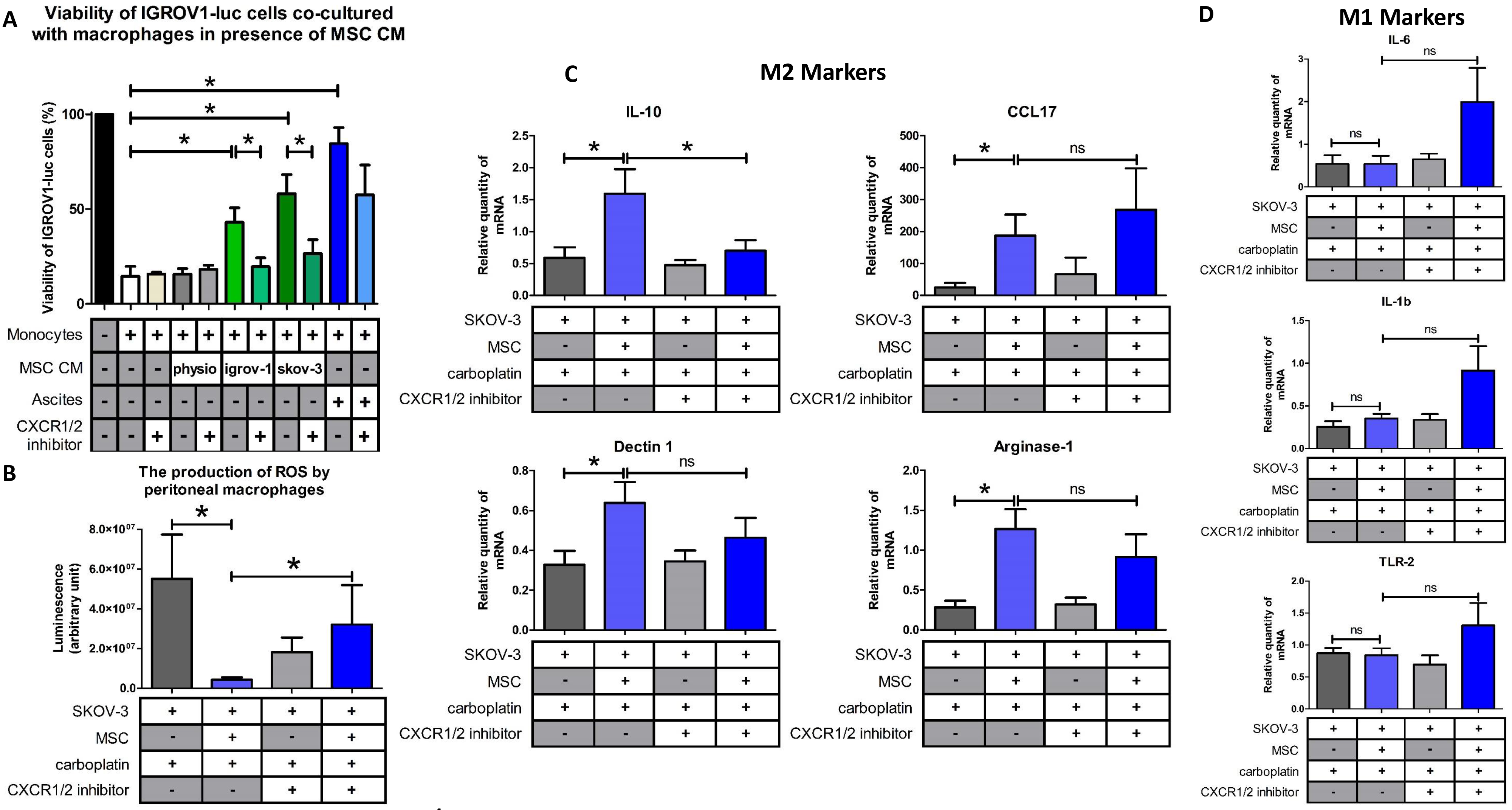
CXCR1/2 inhibition prevented the M2 macrophage polarization induced by CA-MSCs. A: Bar graph represents the % of IGROV-1 cells viability monitored by measuring luciferase activity (n =4). The cytotoxic activity of the macrophages that have been cultured in different media (CM of CA-MSCs (left) or CM of BM-MSCs (physio) or of iCA-MSCs (MSC/igrov-1 CM or MSC/skov-3 CM), or ascites on the IGROV-1luc cells in the presence of a CXCR1/2 inhibitor was evaluated by measuring luciferase activity. B: The production of ROS by peritoneal macrophages isolated from mice injected with SKOV-3 cells admixed or not with MSCs and treated or not with carboplatin +/− CXCR1/2 inhibitor. C, D: Representative gene expression analysis of peritoneal macrophages isolated from mice that have been injected with SKOV-3 cells admixed or not with MSCs and treated with carboplatin +/− CXCR1/2 inhibitor. RNA was extracted and analyzed by RT-qPCR. (8 mice/group). The comparison between groups was performed using a Wilcoxon-Mann Whitney test (independent non-parametric data). P values of <0.05 (*) indicate a significant difference.

Since treatment with the CXCR1/2 inhibitor restored the anti-tumoral properties of macrophages *in vitro*, we wondered if it could also modify the phenotype of macrophages *in vivo*. We analyzed the phenotype of macrophages isolated from the peritonea of human ovarian tumor (SKOV-3) bearing mice which were injected in the presence or absence of MSCs and treated or not with the CXCR1/2 inhibitor (Figure 4C).

M1 macrophages are characterized by the fact that they produce ROS [42]. We measured ROS production in macrophages isolated from mice that were injected with OTCs with or without MSCs and treated or not with a combination carboplatin and the CXCR1/2 inhibitor. As shown Figure 6B, there was a significant difference between macrophages isolated from mice injected with OTCs alone or injected with OTCs admixed with MSCs in the presence of carboplatin. We observed that the presence of MSCs caused a decrease in the ROS production by peritoneal macrophages. When the mice that were injected with OTCs and MSCs were treated with the CXCR1/2 inhibitor, we observed an increase in the production of ROS by the peritoneal macrophages. This suggests a polarization of the macrophages to an M1 phenotype and a tumoricidal effect. This fact could explain that we noticed that the mice co-injected with SKOV-3 cells and MSCs and treated with carboplatin and the CXCR1/2 inhibitor presented a significant slowdown in tumor progression (Figure 4C).

In the same way, we analyzed the phenotype of peritoneal macrophages from tumor bearing mice treated or not with the CXCR1/2 inhibitor. As depicted in Figure 6C, when the mice were injected with a combination of SKOV-3 cells plus MSCs and then treated with carboplatin, there were a high proportion of IL-10, arginase-1, dectin-1 and CCL17 positive macrophages, suggesting a M2 macrophage polarization compared to the mice injected with SKOV-3 cells alone.

When the mice were co-treated with carboplatin and the CXCR1/2 inhibitor, the macrophages did not express as much IL-10, suggesting a shift in macrophage polarization. Concerning the M1 markers, our results suggest a slight upregulation of the expression of IL-1β, IL-6 and TNF-α in the presence of MSCs and the CXCR1/2 inhibitor (Figure 6D), even if gene expression profiles reveal an M1/M2 mixed-polarization phenotype as has previously be shown by Reinartz *et al.*[40,43] in ovarian cancer.

Next, we determined the concentrations of CXCL1, CXCL2 and IL-8 in the peritoneal ascites of the mice treated with carboplatin. As shown in Figure 7 and in line with our previous results, there was an increase in the concentration of CXCL1, CXCL2 and IL-8 in the peritonea of mice injected with SKOV-3 cells and MSCs. When the SKOV-3/MSC tumor bearing mice were treated with the CXCR1/2 inhibitor a decrease in the amount of the three cytokines was observed, suggesting that a blockade of the CXCR1/2 leads to a reduction in the expression of the cytokines (Figure 7).

**Figure 7:**
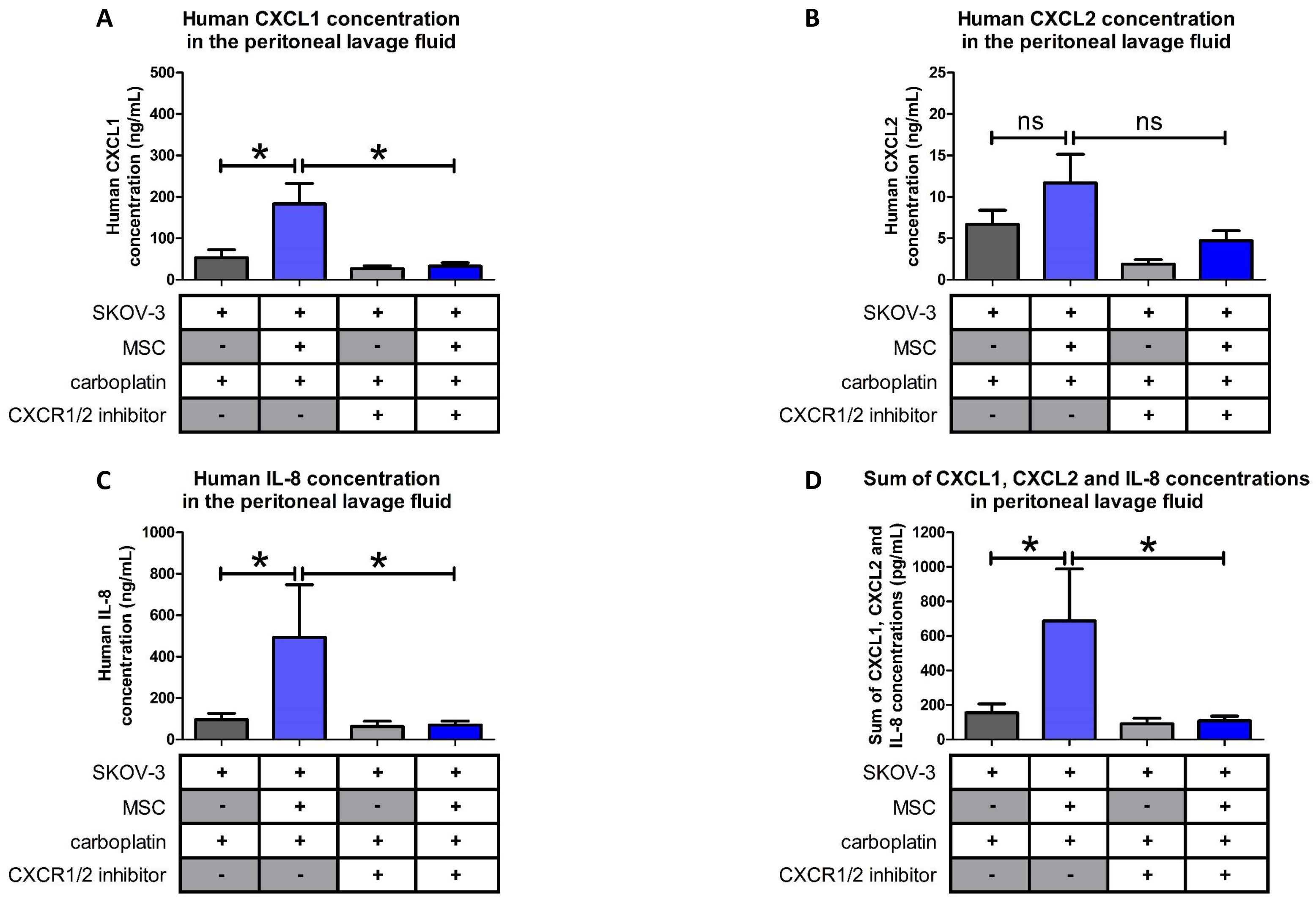
Levels of pro-inflammatory cytokines in the peritonea of mice. Levels of pro-inflammatory cytokines in the peritonea of mice injected with SKOV-3 cells alone or admixed with BM-MSCs and that were treated with carboplatin +/− CXCR1/2 inhibitor. Cytokine levels were evaluated by ELISA (n=8 mice/group). The comparison between groups was performed using a Wilcoxon-Mann Whitney test (independent non-parametric data). P values of <0.05 (*) indicate a significant difference.

All together our results show that through their released factors, CA-MSCs can trigger the differentiation of monocytes to a pro-tumoral M2 phenotype favorable for tumor progression and the acquisition of chemoresistance by OTCs. When CXCR1/2 receptors are inhibited, these CA-MSC-activated macrophages lose their M2 phenotype and show anti-tumoral functions. Inhibition of CXCR1/2 could counteract the pro-tumoral effect of the microenvironment both by sensitizing OTCs to carboplatin and by inducing anti-tumor immunity.

## DISCUSSION

The first aim of our work was to show that MSCs that are recruited to an injury site such as a cancer nodule (CA-MSCs) present some original characteristics compared to naïve BM-MSCs. We showed here that CA-MSCs did not influence the dissemination or proliferation of OTCs either *in vitro* or *in vivo*, but they did play a role in the acquisition of chemoresistance by cancer cells against carboplatin. In addition, we showed that CA-MSCs are involved in the recruitment and polarization of macrophages. Here we characterized precisely the mechanisms by which CA-MSCs exert their biological functions, in particular the role played by the cytokines produced by these cells (CXCR1/2 ligands). We studied the role played by these ligands, while several other authors have mainly focused their attentions on IL-6 and its role in cancer cell proliferation[17,44,45].

CA-MSCs may differentiate from MSCs of different origins. For this study we obtained iCA-MSCs by culturing naïve BM–MSCs from healthy female donors between 60 and 70 years of age in the presence of CM from OTCs or in the presence of ascites. Our aim was to show that we were able to transform naïve BM-MSCs to CA-MSCs through secreted factors present in their microenvironment. We selected a culture period of 21 days as work published by Spaeth et al. showed that the phenotype of MSCs was only modified after they were cultured in the presence of OTC (SKOV-3 cells) CM for more than two weeks[46]. We tried to culture BM-MSCs in the presence of CM from OTCs for a shorter time period (8 or 14 days), but we were unable to convert BM-MSCs to CA-MSCs (iCA-MSCs). This is surprising considering that we could induce the NF-◻B pathway in BM-MSCs with a short exposure (10 min) to the CM (Figure 2B). Ascitic fluid from cancer patients had more or less the same effect on the differentiation of BM-MSCs. These data indicate that factors secreted by tumor cells are sufficient to induce the chemoprotective phenotype of CA-MSCs and that ascites that contain factors secreted by various cell types whose tumor cells result in a comparable phenotype.This work is supported by the recent published data by Coffman et al. who showed that ovarian CA-MSCs arise from tumor mediated reprograming of local tissue MSCs [47].

We evaluated the ability of CM from other epithelial cancer cells, such as pulmonary cancer cells (A549), to activate BM-MSCs and observed the same effect on differentiation as with CM from epithelial OTCs (IGROV-1 or SKOV-3 cells). We found that BM-MSCs cultured in IGROV-1 CM (iCA-MSCs) lost their ability to differentiate into osteoblasts and adipocytes and behaved like CA-MSCs isolated from tumor biopsies of ovarian adenocarcinomas. This work validates the hypothesis that tumor cells can influence the function of MSCs. Our work has shown that through their secreted factors, cancer cells can educate MSCs to the tumor environment whereupon they can educate/reprogram them.[48]. However, we still do not know into what cell type (what particular markers? what function?) these CA-MSCs would have differentiated into. A plausible hypothesis is that they would go on to differentiate into fibroblasts as they express vimentin and PDGFR, but the fact that we could not evidence an important expression of αSMA nor FAP invalidates this hypothesis.

Even if iCA-MSCs obtained from BM-MSCs adopt a phenotype similar to the CA-MSCs found in ovarian adenocarcinomas, it does not prove that CA-MSCs originate from BM-MSCs. These CA-MSCs could originate from MSCs recruited to the tumor site. Nevertheless, they could also derive from “resident” MSCs located to the tissues where the tumor develops. Indeed, MSCs are not restricted to the bone marrow and are found in virtually all tissues, including ovaries [29]. Another possible origin could be adipose tissue, which is a source of MSCs called adipose tissue-derived stromal cells (ADSC). In addition, the epiploon (or omentum) is a double layer of adipose tissue that covers and supports the organs in the lower abdomen. Thus, especially during peritoneal carcinomatosis, ADSCs from the epiploon could constitute a source of CA-MSCs, given their location, close to the OTCs disseminated in the peritoneum.

We found that one hundred genes were upregulated or downregulated in iCA-MSCs compared to their parental BM-MSCs. We analyzed six pairs of cells and obtained reproducible data. Genes were classified based on their implication in angiogenesis, metastasis, tumor cell proliferation, *etc*. We focused on genes implicated in the acquisition of chemoresistance and the recruitment of immune cells. Thus, CXCL1, CXCL2, CXCL3, CXCL5, CXCL6, CXCL8 (IL-8) and CCL5 were found to be upregulated in iCA-MSCs. As CXCL1, CXCL2 and IL-8 interact with the same receptors (CXCR1/2) we analyzed their combined interactions and implications for the acquisition of chemoresistance and the recruitment of immune cells in our model. We confirmed that their upregulation was correlated to patient prognosis. We found that the serum concentrations of these three chemokines, measured in ovarian adenocarcinoma patients at the time of diagnosis, predicted the sensitivity profile of the patients to platinum-based chemotherapy. This was a retrospective study, in the future we would like to perform a prospective study to show that chemokine assays can predict the future resistance of patients to treatment. If this were the case, we could adjust the treatment regimen accordingly.

We found IL-8 to be the most highly upregulated cytokine in CA-MSCs or iCA-MSCs. It has been reported to promote angiogenesis and cancer growth (47), therefore in previous experiments we tried to abolish the acquisition of chemoresistance by tumor cells using an antibody directed against IL-8 [50]. We observed that the antibody had only a very weak effect, suggesting that IL-8 was involved in the acquisition of chemoresistance in association with other molecules and that the pathway involving IL-8 receptors was not inhibited because other molecules could activate these receptors like CXCL1 and CXCL2

Hence, in this study we analyzed the effect of the inhibition of the IL-8 receptors, CXCR1 and CXCR2. Reparixin, a CXCR1/2 inhibitor, has been already used in a mouse model of pancreatic ductal adenocarcinoma to disrupt tumor/fibroblast interactions and improve the survival of tumor bearing mice [32]. We found that the CXCR1/2 inhibitor could reverse the acquisition of chemoresistance by tumor cells both *in vitro* and *in vivo*, in our xenograft model (human SKOV-3 cells injected into nude mice). Therefore, reparixin is very effective at preventing the acquisition of chemoresistance associated with carboplatin treatment. Reparixin is already being tested in clinical trials. Indeed, this drug is being tested in patients with metastatic non-human epidermal growth factor receptor (HER2)-amplified breast cancers in an open label Phase 1b clinical study (REP0111) in combination with a fixed dose of weekly paclitaxel. This study has demonstrated the safety and tolerability of the combination and recorded objective responses. A 30% response rate was recorded, with durable responses of >12 months in two patients [51](https://clinicaltrials.gov/ct2/show/NCT02370238).

The chemokine IL-8 has also been shown to play a key role in the recruitment and activation of polymorphonuclear neutrophils in post-ischemia reperfusion injury after organ transplantation. Hence, reparixin is also in clinical development as a potential key component in an integrated approach for the immunomodulation and control of non-specific inflammatory events. In the case of cancer, IL-8 (as well as CXCL1 or CXCL2) is known to be involved in angiogenesis and the recruitment and activation of immune cells.

Castells *et al.* observed an increased recruitment of macrophages to the tumor site in the presence of CA-MSCs[37]. In fact, monocytes, upon sensing several environmental stresses are recruited to damaged and infected tissues as well as tumor sites and differentiate to macrophages en-route[36]. Several authors report that MSCs facilitate the monocyte to macrophage transition. These TAMs may be of an M1 anti-tumoral phenotype, or of an M2 pro-tumoral phenotype. Previous work performed in our laboratory suggests that CA-MSCs may be able to influence the phenotype of peritoneal macrophages by polarizing them to a pro-tumoral phenotype. In addition, in the case of inflammation, Vasandan *et al.* showed that MSCs induce distinct alterations in human macrophage polarization programs depending on the activation module at macrophage interface[36]. In hematological diseases such as multiple myeloma, Asimakopoulos *et al.* have shown that MSCs and macrophages interact to induce a distinct state of macrophage polarization[52,53]. IL-8, known to induce the chemotaxis of immune cells to the tumor site, may play a role in CA-MSC-induced macrophage recruitment. It could also be the factor responsible for the CA-MSC-induced polarization of macrophages to a pro-tumoral phenotype. Our experimental model allowed us to isolate peritoneal macrophages and determine their phenotype. Dijkgraaf *et al.* have shown that carboplatin chemotherapy increases the number of cancer-supporting M2 macrophages[54]. In our study, we observed that MSCs could induce M2 polarization. In our model, reparixin, when co-administered with carboplatin, decreases the transcription of IL-10, a major marker of M2-type macrophages, and is responsible for an increase in ROS production by peritoneal macrophages and associated to an increase of the transcription M1 markers, as IL-1β, IL-6 and TNF-α.. Therefore, reparixin could sensitize OTCs to carboplatin by decreasing the proportion of type M2 macrophages, or by repolarizing these macrophages to an anti-tumoral phenotype.

In our final experiment when SKOV-3/MSC tumor bearing mice were treated with reparixin, we observed a decrease in the amount of CXCR1/2 ligands in the peritonea, suggesting that a blockade of the CXCR1/2 leads to a decrease in the expression of its ligands. In addition, we suggest that a decrease in the number of tumor cells could reduce the number of MSCs recruited to peritoneal tumors, resulting in a decreased concentration of the chemokines in the peritoneum. The use of chemokine receptor inhibitors could therefore have a direct effect on macrophages by restoring their anti-tumoral activity, moreover they could also have an indirect effect by decreasing the amount of chemokines secreted by CA-MSCs, which would reduce the polarization of macrophages to the M2 phenotype.

In this study, we show that CA-MSCs, which are part of the ovarian tumor microenvironment, can induce tumor cells to become resistant to chemotherapy such as carboplatin. These CA-MSCs secrete chemokines, including IL-8, CXCL1 and CXCL2 that bind to CXCR1/2. The combination of a CXCR1/2 antagonist with platinum-based chemotherapy may be a useful strategy to restore carboplatin sensitivity in OTCs, in addition it may also be able to modify the phenotype of TAMs and reinstate their anti-tumoral activity.

## MATERIALS AND METHODS

### A) Cell culture

The human ovarian adenocarcinoma IGROV-1 (a gift from the Gustave Roussy Institute, Paris) and SKOV-3 (ATCC: HTB-77) cell lines were grown in complete RPMI (RPMI supplemented with fetal calf serum (10%), L-Glutamine (1%) and penicillin / streptomycin (1%)).

Primary BM-MSCs from donors that had undergone orthopedic surgery (Médipole Clinic - Toulouse), CA-MSCs from ovarian cancer patients, were grown in complete DMEM (DMEM supplemented with fetal calf serum (10%), L-Glutamine (1%) and penicillin / streptomycin (1%)).

Primary human macrophages derived from peripheral blood mononuclear cells (PBMC) were cultured in a serum-free medium, the Macrophage-SFM medium (Gibco ™).

All cells were cultured at 37°C in 5% CO_2_. They were regularly treated with Normocin ™ (Invivogen) (100 μg/mL), to prevent mycoplasma contamination.

### B) Ascites and CM

Samples of ascites (n=10) from ovarian cancer patients were obtained from the biological resource center bank at the IUCT-Oncopole. The ascitic fluid was centrifuged at 300 g for 5 minutes. The supernatant was pooled and filtered at 0.2 μm. The CM from BM-MSCs, CA-MSCs, IGROV-1 and SKOV-3 cells corresponded to the supernatant of the cell culture media from confluent cells after 3 days of growth and filtered at 0.2 μm.

### C) Isolation of CA-MSCs from patient biopsies

From fresh tumor biopsies of patients with ovarian cancer, cells were isolated according to their plastic adhesion and then sorted by FACS using the MSC Phenotyping Kit, human (Miltenyi Biotec). CD73+, CD90+, CD105+ and CD14−, CD20−, CD34−, CD45− cells were considered to be CA-MSCs.

### D) Generation of iCA-MSCs

BM-MSCs were cultured in DMEM diluted 1:1 either with CM from the different types of OTCs or ascites. The medium was renewed twice a week. iCA-MSCs were generated after 21 days. Three types of iCA-MSCs: MSC/igrov-1, MSC/skov-3 and MSC/ascite, were induced using the CM of IGROV-1 or SKOV-3 cells or ascites, respectively. To generate CM from the iCA-MSCs, their medium was replaced by complete DMEM and three days later, the various supernatants were filtered at 0.2μm. MSCs cultured in complete DMEM were named physiological BM-MSCs.

### E) Cell viability tests

#### 1) Experiments to analyze the chemosensitivity of IGROV-1 cells

5 × 10^3^ IGROV-1 cells were added per well into 96-well plates in the presence of CM from the different types of MSCs (described above) diluted 1:1 in complete RPMI. After 24 hours, these cells were treated with a range of concentrations of carboplatin (Fresenius Kabi) (15.625 to 1000 μM) with or without the CXCR1/2 inhibitor (AS-62401, AnaSpec, 100 μM). After 48 hours of treatment, cell viability was evaluated using the WST-8 Cell Counting Kit (Dojindo) according to the manufacturer’s instructions.

#### 2) Experiments to assess how macrophage co-culture effects OTC viability

5 × 10^4^ PBMC-derived monocytes were added per well into 96-well white plates and were cultured in CM (diluted 1:1 with Macrophage-SFM (Gibco ™)) from BM-MSCs, CA-MSCs, iCA-MSCs or ascites. After 24 hours, 2.5 × 10^4^ IGROV-1luc were added into the wells. After three days of co-culture, the viability of the IGROV-1luc cells was evaluated by bioluminescence using the Steady-Glo^®^ Luciferase Assay System (Promega) according to the manufacturer’s instructions.

### F) Western blot analysis

Protein extractions were performed and 15 μg of extracted proteins were separated by SDS-PAGE and revealed by antibodies directed against Akt (1: 1000, Cell Signaling # 4691S), p-Akt (1: 1000, Cell Signaling # 4060S), Erk (1: 1000, Cell Signaling # 4696), p-Erk ( 1: 1000; Cell Signaling # 4377S), p-NF-κB p65 (Ser536) (1: 1000; Cell signaling # 3033), NF-κB p65 (1: 1000, Cell signaling # 8242), CXCR1 (1:1000, Sigma-Aldrich SAB2700216), CXCR2 (1: 500, Abcam ab14935), tubulin (1: 1000, Cell signaling # 212146) and actin (1: 1000, Cell signaling # 4968). The intensity of the bands was quantified using ImageJ software.

### G) Transcriptional analysis

#### 1) RNA extractions

RNA was extracted from MSCs using the RNAprotect Cell Reagent and the RNeasy Plus mini kit (QIAGEN) according to the manufacturer’s instructions.

RNA was extracted from murine macrophages after 2 hours of adhesion in 48-well plates, using the RNAqueous-Micro Total RNA Isolation kit (Thermo Fisher Scientific) according to the manufacturer’s instructions.

#### 2) Transcriptomic analysis by Nanostring Technology

The total RNAs extracted from BM-MSCs and iCA-MSCs were analyzed by Nanostring^®^ technology with the “nCounter PanCancer Immune Profiling Panel” to study the transcription of 770 genes[31]. A selection of 29 housekeeping genes was used to calculate the relative amount of target RNA.

#### 3) Transcriptional analysis by RT-qPCR

Complementary DNA (cDNA) derived from total RNA was synthesized using the Verso cDNA Synthesis Kit (Thermo Fisher Scientific), according to the manufacturer’s instructions. PCR was performed using the LightCycler^®^ 480 SYBR Green I Master (Roche), according to the manufacturer’s instructions.

The primers used are listed supplementary figure 5.

### H) Quantification of CXCL1, CXCL2 and IL-8 concentrations by ELISA

The concentrations of murine and human CXCL1, murine and human CXCL2 and human IL-8 were determined by ELISA using the respective ELISA kits, EK0722, EK0723, EK0452 (Boster Biological Technology), ARG80185 (Arigo biolaboratories) and the IL-8 DuoSet^®^ ELISA Development System (R&D Systems), according to the manufacturer’s instructions.

### I) Transcriptional analysis to study macrophage polarization in the presence of iCA-MSCs CM

1 × 10^5^ PBMC were added per well in 48-well plate wells. Macrophages were selected by adhesion (2 hours) then they were immediately brought into contact with CM from BM-MSCs or iCA-MSCs. 24 hours later, macrophage RNAs were extracted as previously described.

### J) *In vivo* experiments

#### 1) Animals

Four to five week old female Swiss nude athymic mice (Charles River laboratories, France) were housed according to the standards of the Federation of European Laboratory Animal Science Associations. They were included in experiments after one week of quarantine.

#### 2) Model 1: Analysis of the peritoneal cancer index

10^7^ SKOV-3 cells with or without 1 × 10^6^ MSCs were injected intraperitoneally (i.p.) into nude mice. Treatments began at 10 days and consisted of 1 injection of carboplatin (200 μl/mouse of a solution of 4.5 mg/mL diluted in 0.9% NaCl) every 7 days for 3 weeks. Tumor progression was monitored after 4 weeks by determining the peritoneal cancer index as described table 1.

#### 3) Model 2: Protocol

10^7^ SKOV-3luc cells with or without 1 × 10^6^ MSCs were injected intraperitoneally (i.p.) into nude mice. Treatments began at 7 days and consisted of 1 injection of carboplatin (200 μl/mouse of a solution of 4.5 mg/mL diluted in 0.9% NaCl) every 7 days for 3 weeks and 1 injection of reparixin (30 mg/kg diluted in a solution of DMSO/PBS (v/v)) 3 times a week for 3 weeks.

#### 4) Model 2: Bioluminescence imaging and measurements

Mice were anesthetized by inhalation of isoflurane (Abbott) at 3% with 1 L/min flow of oxygen. An injection of luciferin (XenoLight D-Luciferin-K + Salt Bioluminescent Substrate, PerkinElmer) is performed intraperitoneally in each mouse 10 minutes prior to analysis with IVIS^®^ Spectrum in vivo imaging system (PerkinElmer), according to the manufacturer’s instructions. A normal behavior of mice was checked before to being returned to their original housing. Images were analyzed by Living Image Software (PerkinElmer) for evaluation and quantification.

#### 5) Model 2: Organ luminescence

After euthanasia, the peritonea, spleens, livers and diaphragms of the mice were removed. These organs were dissected and placed in 96-well white plates. The peritoneal lavage fluid (150 μL) was also placed in white plates. After the addition of 150 μg/mL luciferin (XenoLight D-Luciferin-K + Salt Bioluminescent Substrate, PerkinElmer), organ luminescence was measured using the EnVision™ Multilabel Plate Reader (PerkinElmer).

#### 6) Isolation of macrophages from the peritoneal lavage fluid

The peritoneal lavage fluid (see above) was centrifuged (300 g for 10 minutes). After lysis of the red blood cells, cells were separated using percoll (GE Healthcare Life Sciences, 17-0891), according to the manufacturer’s instructions. 1 × 10^5^ cells from the macrophage-containing fraction were inoculated in 48-well plates. The macrophages were selected by adhesion (2 hours). Their ROS production as well as their mRNA expression levels were evaluated as described below or previously, respectively.

#### 7) Measurement of ROS production

After 2 hours of adhesion at 37°C and 5% CO_2_, the NADPH oxidase activity of peritoneal macrophages (1 × 10^5^) was measured by chemiluminescence in the presence of 60 μM of a chemiluminogenic probe: luminol (5-amino-2,3-dihydro-1,4-phthalazinedione, Sigma-Aldrich). Chemiluminescence production was analyzed with a luminometer (EnVision™ Multilabel Plate Readers, PerkinElmer). Chemiluminescence values were recorded continuously for 90 minutes.

### K) Statistics

For chemoresistance tests, RT-qPCRs, co-cultures, cytometry data, and *in vivo* test results, the comparison between groups was performed using a Wilcoxon-Mann Whitney test (independent non-parametric data). P values of <0.05 (*) and <0.01 (**) indicate a significant difference. An ANOVA test was applied to compare the six BM-MSC control samples versus the six iCA-MSC (MSC/igrov-1) samples. The p values were corrected for multiple tests using the Benjamini & Hochberg (BH) method.

### L) Study approval

All experiments involving animals were performed in accordance with the relevant European guidelines and regulations. The protocols and the experiments were approved by the Claudius Regaud Institute animal ethics committee (approval number: ICR-2015-06).

Human studies: All human biopsies and sera came from patients who provided written informed consent prior to inclusion in the study. The study has been approved by the IUCT-O (Toulouse University Institute of Cancer-Oncopole)) ethics committee

## ACKNOWLEDGMENTS

The authors would like to acknowledge the animal facilities of the Institut Claudius Regaud and the CRCT, as well as the transcriptomic platform at the CRCT. The authors thank Elisabeth Bellard and the “Toulouse Réseau Imagerie” core IPBS facility (Genotoul, Toulouse, France) for technical support. The authors thank Emeline Sarot, Carine Valle, Marie Tosolini, Manon Farcé, Frédéric Lopez and Pôle Technologique du CRCT – Plateaux Génomique et Transcriptomique, Bioinformatique, Cytométrie, Imagerie, INSERM-UMR1037. The authors thank Hubert Lulka for his help with animal imaging and Pr. H. Avet-Loiseau and J. Guillermet-Guibert for administrative support.

This work was funded by a grant from the French government (IDEX 2012) and the Groupe de recherche de l’Institut Claudius Regaud (GRICR). We are grateful to the associations “Phil-Anthrope” and the Rotary club of Lectoure who provided a special grant for this project.

## AUTHOR CONTRIBUTIONS

BC and AC designed the experiments (concept and design, collection and assembly of data, data analysis and interpretation); ALN, MP designed some experiments, performed experiments and analyzed the data (collection and assembly of data); BT, LL, RM, HL, MG, LL, EM, AM, GF provided help with the experiments (collection and assembly of data); ALN and BC wrote the manuscript; JPD provided critical feedback (intellectual support). All authors reviewed the manuscript. The manuscript has been revised for the English by an independent scientific language editing service (AngloScribe)

